# High occupancy of European leaf-toed gecko *Euleptes europaea* in two island stands of *Eucalyptus sp*.: tree selection, co-occurrence and habitat effect

**DOI:** 10.1101/2023.02.08.527781

**Authors:** Grégory Deso, Pauline Priol, Thierry Reynier, Julien Renet

**Author notes:** Corresponding author: Grégory Deso, AHPAM Maison des associations, 384 Route Caderousse, 84100 Orange, France.

## Abstract

The effective conservation of a species requires a thorough knowledge of its ecology. Long considered to live exclusively in rocky habitats, the European leaf-toed gecko (*Euleptes europaea)* has in fact been observed in vegetated and wooded habitats at several locations throughout its range. The tendency of this species to use these habitats seems to be clearly supported by its prehensile tail characteristic of geckos with arboreal behaviour. To better assess tree occupation by *E. europaea* and other co-occurring geckos, a site-occupancy survey was conducted in 2022 on the testing site of DGA (French MoD the Procurement Agency) of Levant Island (Hyères, France). Two stands of *Eucalyptus sp*. containing 68 trees were selected to monitor. One stand lies in an anthropised context, consisting of scattered woodland and clear ground (stand 1), and the other represents a ‘natural’ forest context with dense ground vegetation (stand 2). The results revealed high occupancy by *E. europaea* in both stands, with an average occupancy probability of 0.57 (CI 0.40–0.72). The Mediterranean house gecko (*Hemidactylus turcicus*) and Moorish gecko (*Tarentola mauritanica*) had an average occupancy probability of 0.28 (CI 0.16–0.44) and 0.07 (CI 0.03–0.16) respectively. In stand 2, *E. europaea* was the only gecko species found, suggesting that it is better adapted to this type of forest habitat, which may represent a refuge for this species. In view of these results, the ecology of this species should be reconsidered and the research broadened by systematically including vegetated and forest habitats.

## INTRODUCTION

The conservation of a wild species requires good knowledge of its life-history traits and ecology in order to put in place appropriate management and protection measures (Soulé & Siberloff 1986). Yet sometimes, part of the ecology of a species can go unnoticed or be obscured by lack of knowledge or by ‘expert’ judgement that overlooks uncertainty (Burgman 2005). This is especially true for cryptic species, which are difficult to detect and have a restricted range, so are the least well studied and the least known (de Lima et al. 2011; Martin et al. 2022).

This is the case for the European leaf-toed gecko (*Euleptes europaea*), a tiny cryptic gecko endemic to the western Mediterranean with a mainly insular distribution (Delaugerre 2004). This gecko is a member of the Sphaerodactylidae (Underwood 1954) family, which includes 12 genera of gekkotans with 230 species of terrestrial or arboreal geckos distributed mainly in the Americas, Africa and Asia, of which only *E. europaea* (Gené 1839) is present in Europe (Uetz 2022). First described in 1839 on the island of Sardinia by Gené (1839) as belonging to the genus *Phyllodactylus* (Gray 1828), 158 years later it was assigned to the genus *Euleptes* (Fitzinger 1843) by Bauer et al. (1997). Some anatomical descriptions indicated that this species has a prehensile tail (Fitzinger 1843; Boulenger 1885), which is uncommon in the Sphaerodactylidae family, but characteristic of geckos with arboreal behaviour (Linkem et al. 2008; Grismer et al. 2021). The function of this prehensile tail to move through vegetation was discussed by Lataste (1877) (“*Phyllodactylus* with a hanging tail, usually curling sideways, and clinging with it to the grasses among which it lives”) and later studied in detail by van Eijsden (1983), who found that the tail of *E. europaea* is used as a climbing organ.

It is possible that the propensity of this species to occupy resource-poor environments, often the case of sparsely vegetated Mediterranean habitats, has created a perception bias. Gené (1839) described it as mainly present under the bark of trees and considered it rare under stones “*Sub arborum cortice sat frequens; rarior sub lapidibus*”, and according to Bruno (1976), on the island of Montecristo it was most often found under the bark of heather (*Erica sp*.) and holm oak (*Quercus ilex*). Two years later, Vanni & Lanza (1978) considered this species a bark-dweller, but with an affinity for rocks. More recently, given the current sparsely wooded nature of the Mediterranean islands and islets largely occupied by this species, it has been considered a rocky habitat species both on islands (Delaugerre 1980; Cheylan 1983; Delaugerre & Cheylan 1992) and on the mainland (Salvidio et al. 2010).

The absence of observations of this gecko in some of the tree species surveyed (e.g. cork oak) certainly contributed to this impression that the use of woody plants by *E. europaea* was marginal (Delaugerre 1980). Yet in the last two decades, several records of the species in trees and plants have been reported by naturalists on the islands of Tino (Italy) (Oneto et al. 2008), Levant (France) (Berg & Berg 2010) and Giglio (Italy), where several observations of this gecko were made under the bark of *Eucalyptus sp*. (Fattorini 2010). Indeed, eucalyptus trees shed a great deal of bark, creating sheltering opportunities on the tree itself (under the bark) and on the ground (in the litter). This shelter is utilised by many wildlife species (Verdade et al. 2020; Vasquez et al. 2021), including arboreal geckos in Australia (e.g. *Christinus marmoratus* and *Strophurus congoo*) that are closely associated with semi-dry eucalyptus forests (Davis 2006; Vanderduys 2016).

These historical and recent records seem to point to gaps in our knowledge of the ecology of *E. europaea* and its morphological adaptation to vegetation. This is an urgent concern as the species is near threatened globally (Cox & Temple 2009) and is in danger of extinction over a significant part of its distribution area in southeastern France (Marchand et al. 2017).

To better assess the arboreal behaviour of *E. europaea* and the relevance of the forest environment for the conservation of the species, a study based on the site-occupancy method was carried out in two stands of *Eucalyptus sp*. on Levant Island off the Mediterranean coast of southeastern France. This allowed the identification of the characteristics of the trees, the presence of other gecko species, and the context in which the eucalyptus stands are located, providing a better understanding of the parameters that influence the occupation of the trees by different gecko species.

## MATERIAL & METHODS

### Study area

The study was carried out on Levant Island (43°01’24.5”N 6°27’29.1”E) in southeastern France (Hyères, Var). This island, formed of gneiss and mica schist, has an area of 996 ha and is one of the three main islands of the Hyères archipelago along with Porquerolles and Port-Cros (Tanazacq 1966; Médail et al. 2013). The vegetation mainly consists of tall and impenetrable maquis dominated in some places by Aleppo pine forest (Pinus halepensis).

This plant formation covers most of the area, but there are also several non-native species, including stands of Eucalyptus sp., probably introduced at the beginning of the last century (Jahandiez 1929). The island is located in a sub-humid thermo-Mediterranean bioclimatic zone (Quézel & Médail 2003).

More than half of the island (the eastern side) is covered by a restricted area, and an inhabited civilian zone lies on the western side. The European leaf-toed gecko (*Euleptes europaea*) and the Mediterranean house gecko (*Hemidactylus turcicus*) are the two species of geckos historically present on the island (Jahandiez 1929; Lantz 1931). However, the appearance of the Moorish gecko (*Tarentola mauritanica*) has been noted from 2010 onwards in the civilian area and since 2019 in the restricted area (Deso et al. 2018, 2020).

### Field data collection

For the site-occupancy survey, two allochthonous stands of eucalyptus (*Eucalyptus sp*.) were selected. The first stand was located in an anthropised context and distributed along tracks or road used by motorised vehicles and around technical buildings (stand 1). Further away from the anthropised area, the second stand was characterised by a compact and continuous forest unit, with dense vegetation on the ground and around the trees (stand 2) (Fig. 1). In stand 1, 30 trees were selected and identified (marked with spray paint) and in stand 2, 38 trees were selected and identified.

**Figure 1:**
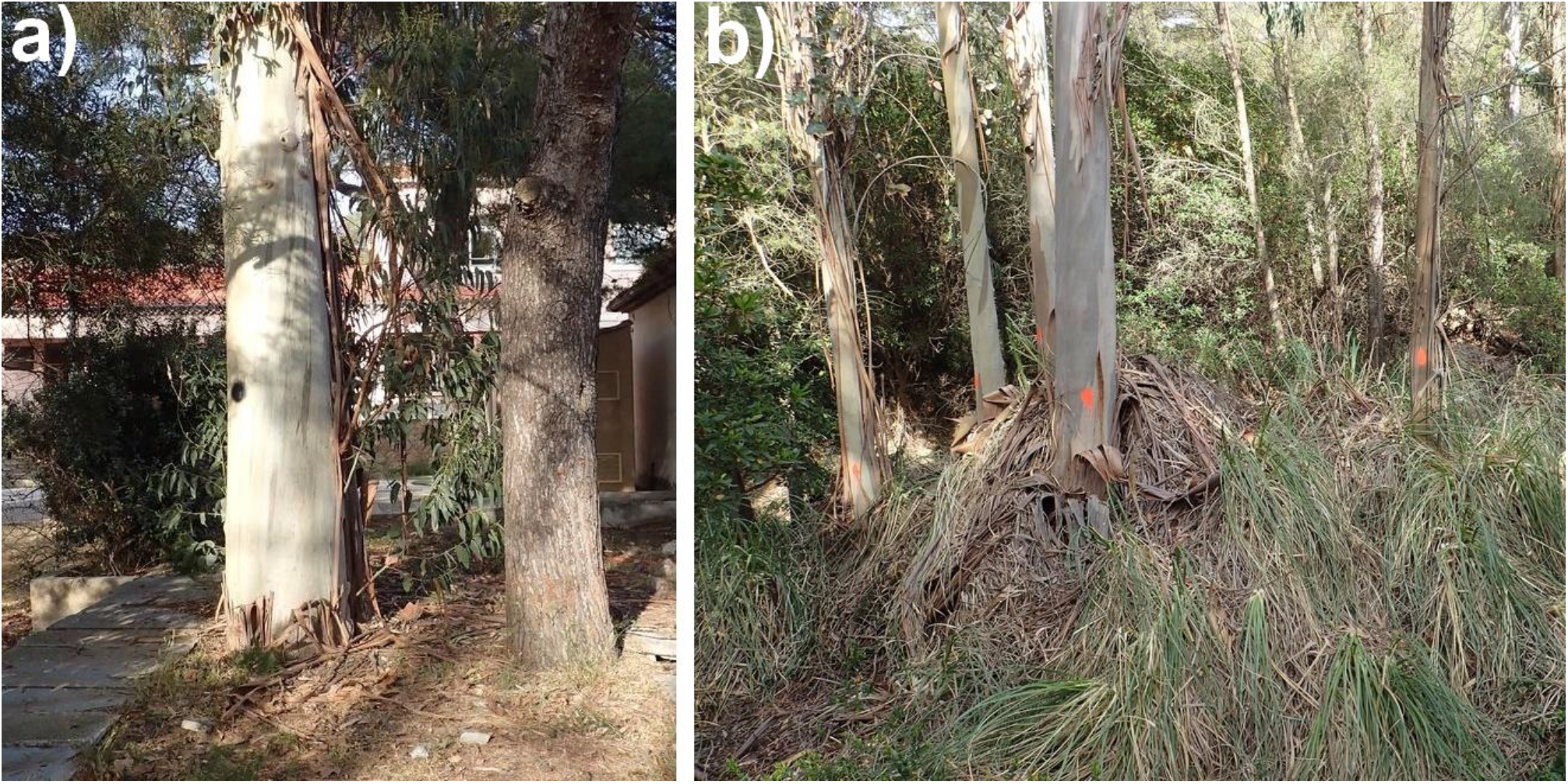
(a) *Eucalyptus sp*. (centre of image) in an anthropised context (stand 1). (b) Dense stand of *Eucalyptus sp*. in a ‘natural’ context (stand 2), with a high degree of ground vegetation cover. (Photos: Julien Renet)

The circumference of all trees was taken using a decameter. The average circumference was 169 cm, ranging from 22 to 550 cm. In stand 1, the average tree circumference was 266 cm (from 120 to 550 cm), and in stand 2 it was 92 cm (from 22 to 423 cm). The average distance between trees was 6.6 m (from 0 to 150 m). The condition of the trees was also noted (alive or dead).

The survey of the trees for *E. europaea* and other co-occurring gecko species, the Mediterranean house gecko (*Hemidactylus turcicus*) and Moorish gecko (*Tarentola mauritanica*), was carried out at night (from 22:00) using headlamps (500 Lumens). The search was carried out under the bark and on the surface of trees up to 2 m above ground level. Each tree was carefully inspected for two minutes. When evident, the gravidity of *E. europaea* females was noted (observation of eggs through the skin).

The survey of the 68 selected trees was replicated three times in the summer (on 01/06/2022, 21/06/2022 and 27/06/2022). One observer carried out the survey for the first session, two for the second session, and three for the last session.

### Statistical analyses

The data collected in the survey was modelled using single-season occupancy models (MacKenzie et al. 2002, 2006). These allowed the estimation of occupation probability, as well as the detection probability of each survey session. We tested the effect of the context – natural or anthropised – (area), the circumference (circumf) and the condition of the tree (state) on the occupancy of each of the three species of geckos found on the island. To take into account the spatial non-independence of the data, an autocorrelation variable (autocor) was created, considering the presence of a species on trees close to the tree concerned (within a 5-m radius). This autocorrelation variable was tested alone and in addition to the other covariates. The numeric circumference covariate was centre-reduced before analysis.

A total of 16 models were fitted for *E. europaea, H. turcicus* and *T. mauritanica*, and maximum likelihood estimates were obtained using the unmarked package (Fiske et al. 2012) in R Studio version 2022.12.0 (R Core team 2022). We assessed the goodness-of-fit of the top-ranked models with the parametric bootstrap using the chi-square as a test statistic with 5000 bootstrap samples. Models were compared using AICc (Burnham & Anderson 2002). We used the entire set of models to draw inferences by computing model-averaged parameter estimates and their unconditional standard errors for the variables appearing in the models with the most support, whereas we model-averaged predictions for the dynamic and detectability parameters from each model (Mazerolle 2013).

## RESULTS

A total of 78 observations were made of geckos on the 68 trees surveyed: 49 *E. europaea*, 23 *H. turcicus* and 6 *T. mauritanica*. In the first session, *E. europaea* was observed 10 times, in the second session 28 times, and in the third session 11 times (Fig. 2). *H. turcicus* was observed 6 times in the first and third sessions, and 11 times in the second. *T. mauritanica* was not observed in the first session, observed 5 times in the second, and once in the third.

**Figure 2:**
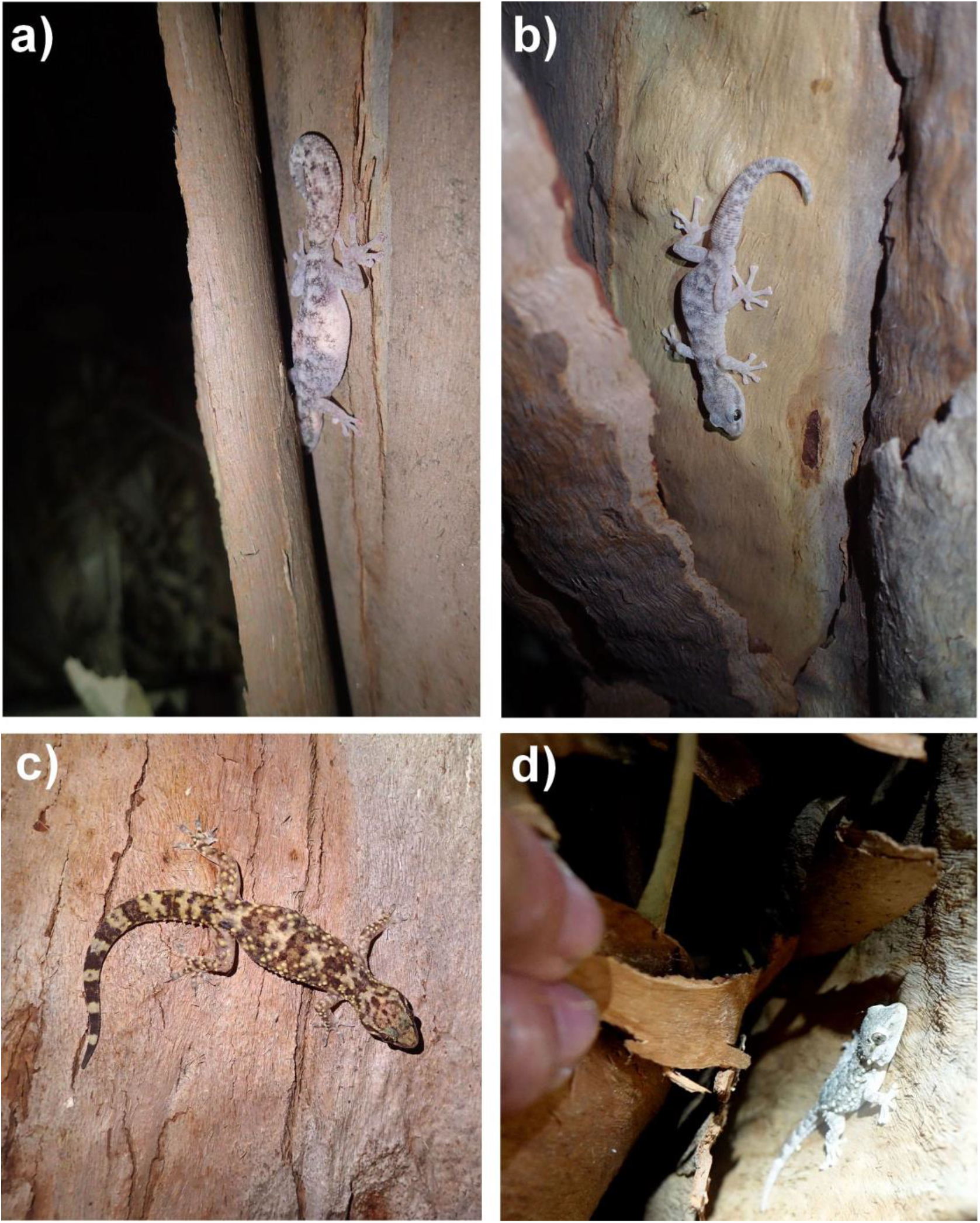
(a) A gravid *E. europaea* female (an egg is visible on the gecko’s left side) takes refuge under the bark of a *Eucalyptus sp*. tree in stand 2. (b) An adult *E. europaea* male in a *Eucalyptus sp*. tree in the anthropised area (stand 1). (c) An adult *H. turcicus* moving on the surface of a *Eucalyptus sp*. tree (stand 1). (d) A subadult *T. mauritanica* under the bark of a *Eucalyptus sp*. tree. (Photos: a, b, c Julien Renet and d, Grégory Deso).

Of the 38 trees in the ‘natural’ area (stand 2), *E. europaea* was found on 19. Of the 30 trees in the anthropised area (stand 1), geckos were found on 23. Of these, 13 trees had only one species (*E. europaea* on 6 trees, *H. turcicus* on 6, and *T. mauritanica* on 1), 9 trees had two species (*E. europaea* and *H. turcicus* on 6 trees, *H. turcicus* and *T. mauritanica* on 2, and *E. europaea* and *T. mauritanica* on 1) and one tree had all three species. *E. europaea* was detected on 14 trees in the anthropised area, *H. turcicus* on 15, and *T. mauritanica* on 5.

### Single-season occupancy models

#### Euleptes europaea

For *E. europaea*, four models were most supported (ΔAICc<2), with an Akaike weight of 0.27 for the first, and 0.19, 0.18 and 0.11 for the following three, respectively (Supplemental Table S1).

These models considered a strong influence of the tree’s state or circumference and/or spatial autocorrelation on occupancy probability and the influence of the survey session on detection probability. The goodness-of-fit of these models was good (respectively P=0.7, c-hat=0.49; P=0.67, c-hat=0.49; P=0.69, c-hat=0.53; and P=0.75, c-hat=0.5). The model averaging showed that spatial autocorrelation was the only variable that had a significant influence on *E. europaea* occupancy (Table 1). Naïve occupancy was estimated at 0.57 (CI 0.40–0.72), or 0.40 (CI 0.22–0.62) when trees within a 5-m radius had no *E. europaea* presence and 0.69 (CI 0.46–0.86) when *E. europaea* was detected in trees nearby.

**TABLE 1.** Model-averaged parameter estimates for *Euleptes europaea* occupancy (Ψ) and detection (p) probability on Levant Island (France). A 95% unconditional confidence interval excluding 0 indicates that the variable had an effect on a parameter.

| Parameter | Estimate | SE | Lower 95% CL | Upper 95% CL |
| --- | --- | --- | --- | --- |
| Occupancy probability ($\Psi$) | | | | |
| Tree state | -1.7508 | 1.2052 | -4.1129 | 0.6113 |
| Tree circumference | 0.5667 | 0.38299 | -0.1836 | 1.3171 |
| Spatial autocorrelation | 1.26040 | 0.6502 | -0.0140 | 2.5347 |

Detection probability was estimated at 0.27 (CI 0.14–0.44) for the first session, 0.74 (CI 0.52–0.89) for the second session, and 0.29 (CI 0.16–0.46) for the third session.

#### Hemidactylus turcicus

For this species, two models were most supported (ΔAICc<2): the first had an Akaike weight of 0.38 and the second a weight of 0.24 (Supplemental Table S2). These models considered an influence of the context (natural or anthropised) on occupancy probability, but the effect was not significant (model-averaged estimates = 10.4739, SE=29.3294, 95% unconditional confidence interval: -47.0108, 67.9585), and the influence of survey session on detection probability. The influence of spatial autocorrelation was not significant (model-averaged estimates = -0.0868, SE=1.5318, 95% unconditional confidence interval: -3.0891, 2.9155). The goodness-of-fit of these models was good (respectively P=0.7, c-hat=0.55 and P=0.53, c-hat=0.9). Naïve occupancy probability was estimated at 0.28 (CI 0.16–0.44); more precisely at 0 (CI 0–1) in natural areas and 0.64 (0.34–0.85) in anthropised areas.

Detection probability was estimated at 0.40 (CI 0.23–0.60); more precisely, 0.33 (CI 0.14– 0.60) for the first and third session and 0.60 (CI 0.31–0.84) for the second session.

#### Tarentola mauritanica

For this species, one model was better supported than the others (ΔAICc<2) with an Akaike weight of 0.60. This model considered the influence of tree circumference on occupancy and survey session on detection with a fair goodness-of-fit (P=0.065, c-hat=3.51) (Supplemental Table S3).

The influence of tree circumference was not significant on occupancy probability (model-averaged estimates = 88.3883, SE=156.3184, 95% unconditional confidence interval: -217.9902, 394.7669).

Naïve occupancy was estimated at 0.07 (CI 0.03–0.16) and naïve detection probability at 0.18 (CI 0.03–0.62), or 0 in the first session, 0.30 (CI 0.13–0.55) in the second and 0.07 (CI 0.01–0.34) in the third.

## DISCUSSION

This study is the first to specifically assess the arboreal behaviour of *E. europaea*, demonstrating an occupancy probability of 0.57 (CI 0.40–0.72) in *Eucalyptus sp*. regardless of the type of area considered (stand 1 or 2). For this species, the effects of circumference, tree condition and context (natural or anthropised) were not significant, indicating that these parameters do not influence this gecko’s tree occupancy. Spatial autocorrelation suggests that the species is gregarious and moves easily from tree to tree if distances are short. For the species *H. turcicus* and *T. mauritanica*, these were only detected in the anthropised area (stand 1) and therefore had a lower average occupancy probability (respectively 0.28 and 0.07).

However, considering only the anthropised area, *H. turcicus* occupied most of the trees (occupancy probability of 0.64), most of the time co-occurring with *E. europaea* or alone, and rarely with *T. mauritanica*. The very low occupancy probability of *T. mauritanica*, even in anthropised areas (estimated at 0.07), is probably explained by its recent arrival on Levant Island (Deso et al. 2018) compared to the other two species studied (Jahandiez 1929; Lantz 1931).

For all three gecko species, these occupancy rates obtained in trees are very likely to be underestimated, as the surveys were only conducted on one type of tree (*Eucalyptus sp*.) and on the lower part of the trees accessible at human height. Other tree and shrub species have been reported in the literature as potential habitats for geckos (e.g. holm oak, aleppo pine, heather, etc.), so it would be of interest to survey these in future.

Detection probability was a function of the survey session for all three species. This was estimated around 0.30 for the first and third sessions for *E. europaea* and *H. turcicus*, and 0.74 (CI 0.52–0.89) for *E. europaea* and 0.60 (CI 0.31–0.84) for *H. turcicus* for the second session. Average detection probability was much lower for *T. mauritanica*, estimated at 0.18 (from 0 to 0.30 depending on the session), probably due to a lower population density. This lower detection may also be due to the fact that this wooded habitat is less optimal for *T. mauritanica*.

It should be noted that the effect of the session conflates the detection ability of the observers and any meteorological effects. However, as the number or experience of observers was not greater in session two, it seems safe to presume that the differences were due to fluctuations in gecko activity. In any case, these results highlight the need to make several passes when surveying reptiles in order to take into account detection probability of these often rare and secretive species (Mazerolle et al. 2007). Otherwise, there is a risk of missing their presence and of drawing erroneous conclusions about their ecology.

While our results found that three gecko species use trees in the study site, the exclusive presence of *E. europaea* in stand 2 is revealing. This could indicate that the morphology of this species is most adapted to environments with dense vegetation. Unlike *H. turcicus* and *T. mauritanica, E. europaea* has a prehensile tail bearing adhesive pads with sensory capabilities (Griffing et al. 2021) found in many species of arboreal geckos (Bauer 1998; Bauer & Menegon 2006). From an evolutionary point of view, a tail can act as a fifth limb, facilitating both balance when resting and slow climbing within densely vegetated habitats on the ground and in the upper forest strata (Jusufi et al. 2008). The high vegetation cover observed in stand 2 is likely to represent a barrier for less well-adapted species. Like other members of the Sphaerodactylidae family (e.g. *Coleodactylus natalensis*, De Sousa & Freire 2011) *E. europaea* may also be a more shade-loving species and a passive thermoregulator that is better adapted to the forest environment than *T. mauritanica* and *H. turcicus*.

Eucalyptus stands characterised by dense ground vegetation could thus be used as refuges by *E. europaea*, allowing the gecko to mitigate the effects of potential competitive interactions with co-occurring species, in particular *T. mauritanica*, which was recently introduced on Levant Island (Deso et al. 2018). The latter is a massive gecko known for its strong territoriality and frequent agonistic behaviour (Lisicic et al. 2012; Salvador 2016), and it is suspected of having caused the decline of *E. europaea* in a mainland locality (Renet et al. *in prep*.). The arrival of *T. mauritanica* on Levant Island is likely to lead to behavioural responses in the *E. europaea* population: for example, asynchronous activity rates (Luiselli & Capizzi 1999) or the emergence of avoidance strategies such as retreating into specialised habitats (Delaugerre et al. 2019; Williams et al. 2020a, b). This will certainly have a cost that needs to be assessed in the future. In a mainland site in southern Tuscany (Italy), Radi & Zuffi (2022) found that the effective spatial segregation between *T. mauritanica, H. turcicus* and *E. europaea* suggested a different rhythm of activity of *E. europaea* compared to the other two. The biotic relationships between these gecko species remain poorly known and merit further research, with priority given to newly colonised areas.

Our findings lead us to consider *Eucalyptus sp*. stands as habitats of high importance for *E. europaea* conservation despite the fact that these trees are non-native. The peeling bark sheds in long ribbons, which seem optimal for geckos to find suitable refuges, lay eggs (10% of our *E. europaea* observations concerned gravid females) and search for prey (Fig. 2).

To further the results of this study, it would be valuable to assess occupancy in other tree species such as Aleppo pine (*Pinus halepensis*), tree heather (*Erica arborea*), holm oak (*Quercus ilex*) and cork oak (*Quercus suber*), which are widely distributed on Levant Island. It is also imperative to better understand the use of vegetation by *E. europaea* and how certain forest structures may prohibit colonisation by other gecko species.

In conclusion, we advocate a reconsideration of *E. europaea* ecology that broadens its habitat to include vegetated areas and woodland, as observed in our study and by earlier naturalists. It seems clear that the image we have of *E. europaea* distribution is biased by having carried out surveys mainly in rocky habitats, and that our scope of research for this species should be widened. This argument is reinforced by the recent chance discovery of *E. europaea* in an Aleppo pine forest in the Naples area, about 330 km from the nearest previous continental record of its sighting (Di Nicola et al. 2022).

## ACKNOWLEDGEMENTS

We would like to warmly thank the staff of the testing site of DGA (the French MoD Procurement Agency), Lucile Objois, Céline Monserat, Patrice Ortola, Charly Gicqueau, Laura Vetter and Sandrine Perroni who made this study possible and facilitated the logistics on the study site. We also thank Aloys Crouzet, Stéphanie Cappellano, Paul Simar Bayet and Tony Genova who offered support and helped us in the field, Aaron Bauer (Villanova University, United States) for his useful comments and Elise Bradbury for her work in proofreading the English.

## AUTHOR CONTRIBUTION

JR and PP conceptualised and designed the study. GD, TR and JR supervised the field data collection and the overall study. PP carried out the data analysis. GD, PP and JR wrote the first manuscript draft in equal parts. All authors revised and approved the manuscript.

## SUPPLEMENTAL DATA

Supplemental data for this article can be accessed at:

## SUPPLEMENTAL DATA

**Supplemental Table S1.** 16 occupancy models based on the second-order Akaike information criterion (AICc), showing the distance between each model and the top-ranked model (ΔAICc), Akaike weights (wi) and the number of estimated parameters (K) for *Euleptes europaea* on Levant Island during 2022. Ψ is occupancy probability and p detection probability.

| No. | Models | K | AICc | $\Delta AICc$ | $W_i$ |
| --- | --- | --- | --- | --- | --- |
| 1 | $\Psi(\text{circumf}+\text{autocor}) p(t)$ | 6 | 204.20 | 0.00 | 0.27 |
| 2 | $\Psi(\text{autocor}) p(t)$ | 5 | 204.84 | 0.64 | 0.19 |
| 3 | $\Psi(\text{state}+\text{autocor}) p(t)$ | 6 | 204.96 | 0.76 | 0.18 |
| 4 | $\Psi(\text{state}) p(t)$ | 5 | 205.97 | 1.77 | 0.11 |
| 5 | $\Psi(.) p(t)$ | 4 | 206.55 | 2.36 | 0.08 |
| 6 | $\Psi(\text{Circonf}) p(t)$ | 5 | 206.87 | 2.67 | 0.07 |
| 7 | $\Psi(\text{area}+\text{autocor}) p(t)$ | 6 | 207.16 | 2.96 | 0.06 |
| 8 | $\Psi(\text{area}) p(t)$ | 5 | 208.81 | 4.61 | 0.03 |
| 9 | $\Psi(\text{state}+\text{autocor}) p(.)$ | 4 | 220.94 | 16.74 | 0.00 |
| 10 | $\Psi(\text{circumf}+\text{autocor}) p(.)$ | 4 | 221.17 | 16.97 | 0.00 |
| 11 | $\Psi(.) p(.)$ | 3 | 221.19 | 16.99 | 0.00 |
| 12 | $\Psi(\text{state}) p(.)$ | 3 | 222.31 | 18.12 | 0.00 |
| 13 | $\Psi(.) p(.)$ | 2 | 223.04 | 18.84 | 0.00 |
| 14 | $\Psi(\text{circumf}) p(.)$ | 3 | 223.32 | 19.12 | 0.00 |
| 15 | $\Psi(\text{area}+\text{autocor}) p(.)$ | 4 | 223.45 | 19.25 | 0.00 |
| 16 | $\Psi(\text{area}) p(.)$ | 3 | 225.16 | 20.96 | 0.00 |

**Supplemental Table S2.** 16 occupancy models based on the second-order Akaike information criterion (AICc), showing the distance between each model and the top-ranked model (ΔAICc), Akaike weights (wi) and the number of estimated parameters (K) for *Hemidactylus turcicus* on Levant Island during 2022. Ψ is occupancy probability and p detection probability.

| No. | Models | K | AICc | $\Delta\text{AICc}$ | $w_i$ |
| --- | --- | --- | --- | --- | --- |
| 1 | $\Psi(\text{area}) p(.)$ | 3 | 105.29 | 0.00 | 0.38 |
| 2 | $\Psi(\text{area}) p(t)$ | 5 | 106.21 | 0.92 | 0.24 |
| 3 | $\Psi(\text{area}+\text{autocor}) p(.)$ | 4 | 107.41 | 2.11 | 0.13 |
| 4 | $\Psi(\text{circumf}) p(.)$ | 3 | 108.40 | 3.11 | 0.08 |
| 5 | $\Psi(\text{area}+\text{autocor}) p(t)$ | 6 | 108.48 | 3.18 | 0.08 |
| 6 | $\Psi(\text{Circonf}) p(t)$ | 5 | 109.97 | 4.68 | 0.04 |
| 7 | $\Psi(\text{circumf}+\text{autocor}) p(.)$ | 4 | 110.37 | 5.07 | 0.03 |
| 8 | $\Psi(\text{circumf}+\text{autocor}) p(t)$ | 6 | 112.07 | 6.77 | 0.01 |
| 9 | $\Psi(\text{autocor}) p(.)$ | 3 | 131.80 | 26.51 | 0.00 |
| 10 | $\Psi(\text{autocor}) p(t)$ | 5 | 132.72 | 27.43 | 0.00 |
| 11 | $\Psi(.) p(.)$ | 2 | 133.27 | 27.98 | 0.00 |
| 12 | $\Psi(\text{state}+\text{autocor}) p(.)$ | 4 | 133.36 | 28.07 | 0.00 |
| 13 | $\Psi(.) p(t)$ | 4 | 134.05 | 28.76 | 0.00 |
| 14 | $\Psi(\text{state}+\text{autocor}) p(t)$ | 6 | 134.49 | 29.19 | 0.00 |
| 15 | $\Psi(\text{state}) p(.)$ | 3 | 135.02 | 29.72 | 0.00 |
| 16 | $\Psi(\text{state}) p(t)$ | 5 | 135.94 | 30.65 | 0.00 |

**Supplemental Table S3.** 16 occupancy models based on the second-order Akaike information criterion (AICc), showing the distance between each model and the top-ranked model (ΔAICc), Akaike weights (wi) and the number of estimated parameters (K) for *Tarentola mauritanica* on Levant Island during 2022. Ψ is occupancy probability and p detection probability.

| No. | Models | K | AICc | $\Delta\text{AICc}$ | $w_i$ |
| --- | --- | --- | --- | --- | --- |
| 1 | $\Psi(\text{circumf}) p(t)$ | 5 | 38.43 | 0.00 | 0.60 |
| 2 | $\Psi(\text{area}+\text{autocor}) p(t)$ | 6 | 41.54 | 3.11 | 0.13 |
| 3 | $\Psi(\text{circumf}) p(.)$ | 3 | 42.99 | 4.56 | 0.06 |
| 4 | $\Psi(\text{area}) p(t)$ | 5 | 43.01 | 4.58 | 0.06 |
| 5 | $\Psi(\text{autocor}) p(t)$ | 5 | 43.14 | 4.71 | 0.06 |
| 6 | $\Psi(\text{circumf}+\text{autocor}) p(t)$ | 6 | 44.32 | 5.89 | 0.03 |
| 7 | $\Psi(\text{circumf}+\text{autocor}) p(.)$ | 4 | 45.09 | 6.66 | 0.02 |
| 8 | $\Psi(\text{state}+\text{autocor}) p(t)$ | 6 | 45.13 | 6.70 | 0.02 |
| 9 | $\Psi(\text{area}+\text{autocor}) p(.)$ | 4 | 47.92 | 9.49 | 0.01 |
| 10 | $\Psi(.) p(t)$ | 4 | 49.36 | 10.94 | 0.00 |
| 11 | $\Psi(\text{area}) p(.)$ | 3 | 49.54 | 11.12 | 0.00 |
| 12 | $\Psi(\text{autocor}) p(.)$ | 3 | 49.68 | 11.25 | 0.00 |
| 13 | $\Psi(\text{state}) p(t)$ | 5 | 50.74 | 12.31 | 0.00 |
| 14 | $\Psi(\text{state}+\text{autocor}) p(.)$ | 4 | 51.51 | 13.09 | 0.00 |
| 15 | $\Psi(.) p(.)$ | 2 | 56.04 | 17.61 | 0.00 |
| 16 | $\Psi(\text{state}) p(.)$ | 3 | 57.27 | 18.85 | 0.00 |

