## Supplemental Table for "High occupancy of European leaf-toed gecko *Euleptes europaea* in two island stands of *Eucalyptus sp*.: tree selection, co-occurrence and habitat effect"

Supplemental data

Supplemental Table S1. 16 occupancy models based on the second-order Akaike information criterion (AICc), showing the distance between each model and the top-ranked model (ΔAICc), Akaike weights (wi) and the number of estimated parameters (K) for *Euleptes europaea* on Levant Island during 2022. Ψ is occupancy probability and p detection probability.

| No. | Models | K | AICc | ΔAICc | Wi |
| --- | --- | --- | --- | --- | --- |
| 1 | Ψ(circumf+autocor) p(t) | 6 | 204.20 | 0.00 | 0.27 |
| 2 | Ψ(autocor) p(t) | 5 | 204.84 | 0.64 | 0.19 |
| 3 | Ψ(state+autocor) p(t) | 6 | 204.96 | 0.76 | 0.18 |
| 4 | Ψ(state) p(t) | 5 | 205.97 | 1.77 | 0.11 |
| 5 | Ψ(.) p(t) | 4 | 206.55 | 2.36 | 0.08 |
| 6 | Ψ(Circonf) p(t) | 5 | 206.87 | 2.67 | 0.07 |
| 7 | Ψ(area+autocor) p(t) | 6 | 207.16 | 2.96 | 0.06 |
| 8 | Ψ(area) p(t) | 5 | 208.81 | 4.61 | 0.03 |
| 9 | Ψ(state+autocor) p(.) | 4 | 220.94 | 16.74 | 0.00 |
| 10 | Ψ(circumf+autocor) p(.) | 4 | 221.17 | 16.97 | 0.00 |
| 11 | Ψ(.) p(.) | 3 | 221.19 | 16.99 | 0.00 |
| 12 | Ψ(state) p(.) | 3 | 222.31 | 18.12 | 0.00 |
| 13 | Ψ(.) p(.) | 2 | 223.04 | 18.84 | 0.00 |
| 14 | Ψ(circumf) p(.) | 3 | 223.32 | 19.12 | 0.00 |
| 15 | Ψ(area+autocor) p(.) | 4 | 223.45 | 19.25 | 0.00 |
| 16 | Ψ(area) p(.) | 3 | 225.16 | 20.96 | 0.00 |

Supplemental Table S2. 16 occupancy models based on the second-order Akaike information criterion (AICc), showing the distance between each model and the top-ranked model (ΔAICc), Akaike weights (wi) and the number of estimated parameters (K) for *Hemidactylus turcicus* on Levant Island during 2022. Ψ is occupancy probability and p detection probability.

| No. | Models | K | AICc | ΔAICc | Wi |
| --- | --- | --- | --- | --- | --- |
| 1 | Ψ(area) p(.) | 3 | 105.29 | 0.00 | 0.38 |
| 2 | Ψ(area) p(t) | 5 | 106.21 | 0.92 | 0.24 |
| 3 | Ψ(area+autocor) p(.) | 4 | 107.41 | 2.11 | 0.13 |
| 4 | Ψ(circumf) p(.) | 3 | 108.40 | 3.11 | 0.08 |
| 5 | Ψ(area+autocor) p(t) | 6 | 108.48 | 3.18 | 0.08 |
| 6 | Ψ(Circonf) p(t) | 5 | 109.97 | 4.68 | 0.04 |
| 7 | Ψ(circumf+autocor) p(.) | 4 | 110.37 | 5.07 | 0.03 |
| 8 | Ψ(circumf+autocor) p(t) | 6 | 112.07 | 6.77 | 0.01 |
| 9 | Ψ(autocor) p(.) | 3 | 131.80 | 26.51 | 0.00 |
| 10 | Ψ(autocor) p(t) | 5 | 132.72 | 27.43 | 0.00 |
| 11 | Ψ(.) p(.) | 2 | 133.27 | 27.98 | 0.00 |
| 12 | Ψ(state+autocor) p(.) | 4 | 133.36 | 28.07 | 0.00 |
| 13 | Ψ(.) p(t) | 4 | 134.05 | 28.76 | 0.00 |
| 14 | Ψ(state+autocor) p(t) | 6 | 134.49 | 29.19 | 0.00 |
| 15 | Ψ(state) p(.) | 3 | 135.02 | 29.72 | 0.00 |
| 16 | Ψ(state) p(t) | 5 | 135.94 | 30.65 | 0.00 |

Supplemental Table S3. 16 occupancy models based on the second-order Akaike information criterion (AICc), showing the distance between each model and the top-ranked model (ΔAICc), Akaike weights (wi) and the number of estimated parameters (K) for *Tarentola mauritanica* on Levant Island during 2022. Ψ is occupancy probability and p detection probability.

| No. | Models | K | AICc | ΔAICc | Wi |
| --- | --- | --- | --- | --- | --- |
| 1 | Ψ(circumf) p(t) | 5 | 38.43 | 0.00 | 0.60 |
| 2 | Ψ(area+autocor) p(t) | 6 | 41.54 | 3.11 | 0.13 |
| 3 | Ψ(circumf) p(.) | 3 | 42.99 | 4.56 | 0.06 |
| 4 | Ψ(area) p(t) | 5 | 43.01 | 4.58 | 0.06 |
| 5 | Ψ(autocor) p(t) | 5 | 43.14 | 4.71 | 0.06 |
| 6 | Ψ(circumf+autocor) p(t) | 6 | 44.32 | 5.89 | 0.03 |
| 7 | Ψ(circumf+autocor) p(.) | 4 | 45.09 | 6.66 | 0.02 |
| 8 | Ψ(state+autocor) p(t) | 6 | 45.13 | 6.70 | 0.02 |
| 9 | Ψ(area+autocor) p(.) | 4 | 47.92 | 9.49 | 0.01 |
| 10 | Ψ(.) p(t) | 4 | 49.36 | 10.94 | 0.00 |
| 11 | Ψ(area) p(.) | 3 | 49.54 | 11.12 | 0.00 |
| 12 | Ψ(autocor) p(.) | 3 | 49.68 | 11.25 | 0.00 |
| 13 | Ψ(state) p(t) | 5 | 50.74 | 12.31 | 0.00 |
| 14 | Ψ(state+autocor) p(.) | 4 | 51.51 | 13.09 | 0.00 |
| 15 | Ψ(.) p(.) | 2 | 56.04 | 17.61 | 0.00 |
| 16 | Ψ(state) p(.) | 3 | 57.27 | 18.85 | 0.00 |
